# Genomic glucocorticoid receptor effects guide acute stress-induced delayed anxiety and basolateral amygdala spine plasticity in rats

**DOI:** 10.1101/2022.09.19.508565

**Authors:** Leonardo S. Novaes, Leticia M. Bueno-de-Camargo, Amadeu Shigeo-de-Almeida, Vitor A. L. Juliano, Ki Goosens, Carolina D. Munhoz

## Abstract

Anxiety, a state related to anticipatory fear, can be adaptive in the face of environmental threats or stressors. However, anxiety can also become persistent and manifest as anxiety-and stress-related disorders, such as generalized anxiety or post-traumatic stress disorder (PTSD). In rodents, systemic administration of glucocorticoids (GCs) or short-term restraint stress induces anxiety-like behaviors and dendritic branching within the basolateral complex of the amygdala (BLA) ten days later. Additionally, increased arousal-related memory retention mediated by elevated GCs requires concomitant noradrenaline (NE) signaling, both acting in the BLA. It is unknown whether GCs and NE play a role in the delayed acute stress-induced effects on behavior and BLA dendritic plasticity. Here, inhibiting corticosterone (CORT) elevation during two hours of restraint stress prevents stress-induced increases in delayed anxiety-like behavior and BLA dendritic spine density in rats. Also, we show that the delayed acute stress-induced effects on behavior and morphological alterations are critically dependent on genomic glucocorticoid receptor (GR) actions in the BLA. Unlike CORT, the pharmacological enhancement of NE signaling in the BLA was insufficient to drive delayed anxiety-related behavior. Nonetheless, the delayed anxiety-like behavior ten days after acute stress requires NE signaling in the BLA during stress exposure. Therefore, we define the essential roles of two stress-related hormones for the late stress consequences, acting at two separate times: CORT, via GR, immediately during stress and NE, via beta-adrenoceptors, during the expression of delayed anxiety.

**Significance Statement:** The dysregulation in orchestrating and finetuning major stress-related neural circuitries leads to enhanced reactivity and other altered ways of coping with threatening situations, predisposing humans to multiple psychiatric disorders, including anxiety and PTSD. Given the tremendous burden of affective disorders, we must advance our understanding of stress neurobiology and translate this into improved treatments. Here we showed that the absence of neuronal genomic GR signaling in the BLA prevented delayed effects on anxiety-like behavior and dendritic spine density ten days after stressor exposure. We also demonstrate that CORT, via GR and immediately at stress and NE, via beta-adrenoceptors, during the expression of delayed behavior contribute to the late stress consequences.

## Introduction

The body’s physiological and behavioral responses to stressful stimuli have critical adaptive value as it prepares to deal with threatening conditions. The almost immediate release of norepinephrine (NE) in the brain during emotional arousal binds to alpha and beta-adrenergic receptors (1). Tens of minutes later, the release of glucocorticoid hormones (GCs) peaks, driving many acute adaptations to stress through actions at glucocorticoid receptors (GRs) [for a review, (2)]. A seminal study showed that elevated corticosterone (CORT; the principal GC in rodents) levels enhance memory retention in tasks with high arousal content (3). The authors also demonstrated that arousal is determined by the NE levels that reach the amygdala’s basolateral complex (BLA) (3).

However, maladaptation can sometimes follow stressor exposure (4). Exacerbated fear responses to stimuli that would not usually elicit such a response, avoidance behavior, and persistent anxiety symptoms are characteristic of anxiety- and stress-related disorders, such as generalized anxiety disorder and post-traumatic stress disorder (PTSD) (5). The relationship between an acute stressor and the delayed manifestation of anxiety symptoms has been widely studied, mainly because it resembles stress-related psychiatric disorders like PTSD.

Several studies have shown that a single two-hour stress session can produce anxiety-like behavior ten days later (6-9). In addition to promoting delayed anxiety-related behavior in rats, two-hour restraint stress and single prolonged stress protocols are associated with deficits in fear extinction memory (10, 11). Some studies have shown that acute two-hour immobilization stress or systemic CORT elevation promotes anxiety-like behavior in rats 10- 12 days later, accompanied by increased spine density and dendritic arborization in the BLA (7, 12, 13). This delayed effect of stress on behavior is associated with morphological changes in the BLA, suggesting that increased synaptic contacts in this brain area are the basis for the increased anxiogenic drive. Thus, since hyperexcitability of the amygdala has also been linked to anxiety (14), it is plausible that BLA dendritic tree hypertrophy plays a causal role in anxiety.

Although there is a clear relationship between a transient increase in CORT levels and emergent anxiety-related behavior, it is unclear whether CORT acts directly on BLA neurons contributing to the behavioral and morphological effects after an acute stress episode. It also remains to be clarified whether these morphological changes in the BLA are sufficient to promote anxiety or if there is a dependence on another mediator, in addition to CORT, for the delayed behavioral manifestation associated with acute stress. Thus, the present study sought to evaluate the role of GR, particularly the genomic actions of this receptor, in the BLA at the time of stress in stress-induced behavioral and morphological outcomes. Considering the pivotal participation of NE in modulating emotional arousal and neuronal excitability (15), we also explored the role of BLA adrenergic receptors during stress-induced anxiety-related behavior development.

## Results

### Systemic CORT release is necessary for the delayed stress-induced anxiety-like behavior and increased BLA dendritic spine density

To confirm whether the acute elevation of CORT plasma levels is essential to the stress-related behavioral and neuroplastic effects, we inhibited the stress-induced rise in this hormone’s blood levels by administering metyrapone (Met). This compound inhibits the synthesis of GCs, before stress exposure. The behavioral outcomes were then assessed ten days later (Fig. 1A). As shown in Fig. 1B, pre-stress Met administration effectively prevented the acute stress-induced increase in CORT plasma levels measured immediately after stress exposure. In contrast, stressed animals administered saline presented higher plasma CORT concentrations than non-stressed animals (P < 0.001) and stressed animals treated with Met (P < 0.001).

**Figure 1.**
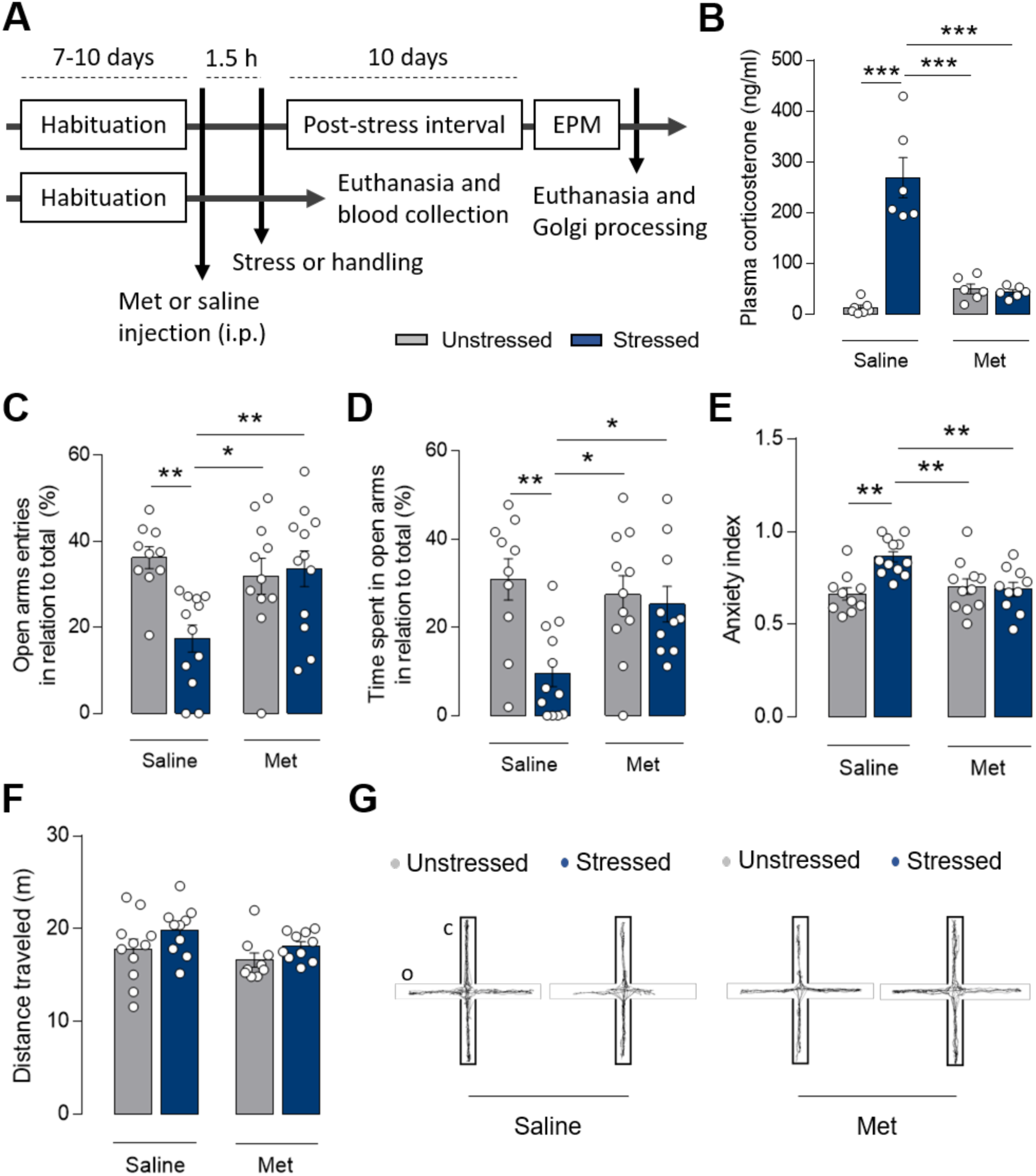
CORT synthesis inhibition during stress prevented the delayed emergence of anxiety-like behavior. (A) Schematic representation of the experimental design showing the timeline of vehicle (saline) or metyrapone (Met) injections, the stress submission, and the EPM behavioral test. One cohort of animals was euthanized immediately after the end of the stress session. (B) Previous administration of Met prevented the stress-induced increase in CORT plasma levels measured immediately after stress (n = 6–7 animals for each group, two-way ANOVA: stress F1, 21 = 40.34, P < 0.0001; Met F1, 21 = 22.81, P = 0.0001; stress-Met interaction F1, 21 = 44.35, P < 0.0001). (C and D) Stressed animals treated with saline exhibited reduced open arm entries and time spent in the EPM test compared to saline-treated non-stressed animals and stressed and non-stressed Met-treated animals (C; n = 10– 12 animals for each group, two-way ANOVA: stress F1, 40 = 4.692, P = 0.0363; Met F1, 40 = 3.23, P = 0.0799; stress-Met interaction F1, 40 = 8.9, P = 0.0048; D; n = 10–12 animals for each group, two-way ANOVA: stress F1, 39 = 8.712, P = 0.0053; Met F1, 39 = 5.783, P = 0.0210; stress-Met interaction F1, 39 = 2.352, P = 0.1332). (E) Stress increased the anxiety index in saline-treated but not in Met-treated rats (n = 10–12 animals for each group, two-way ANOVA: stress F1, 39 = 6.85, P = 0.0126; Met F1, 39 = 3.786, P = 0.0589; stress-Met interaction F1, 39 = 9.282, P = 0.0041). (F) Neither stress nor Met administration influenced the animal’s locomotor activity in the EPM (n = 9–12 animals for each group, two-way ANOVA: stress F1, 34 = 2.388, P = 0.1315; Met F1, 34 = 3.78, P = 0.0602; stress-Met interaction F1, 34 = 0.1839, P = 0.6707). (G) Representative tracking plots for the average motion of each experimental group in the EPM. Results are represented as the mean ± SEM. Tukey’s multiple comparisons post-hoc test. Significance differences between groups are indicated as * P < 0.05; ** P < 0.01; and *** P < 0.001.

Ten days after restraint stress, we assessed the anxiety-like behavior using the elevated plus-maze (EPM) test. Like the previous results, pre-stress Met administration blunted the acute stress-induced anxiety-like behavior onset and development. Indeed, Met-administered stressed animals presented higher open arm exploration than saline-treated stressed animals (P < 0.01, Fig. 1C; P < 0.05, Fig. 1D) and the same exploration profile as the saline non-stressed animals (Fig. 1C and D). The anxiety index confirmed these data, being higher in saline-treated stressed animals than in non-stressed (P < 0.01) and Met-treated stressed animals (P < 0.01) (Fig. 1E). Finally, neither stress nor Met treatment influenced the animals’ motor behavior, as indicated by their total displacement in the maze (Fig. 1F and G).

We analyzed the effects of pre-stress Met administration on the delayed stress-induced increase in dendritic spine density in the BLA using Golgi staining in a subset of the rats that underwent behavioral evaluation (n = 4/group). We confirmed that a single two-hour restraint stress exposure augmented the spine density in the primary (P < 0.001, Fig. 2A), secondary (P < 0.001, Fig. 2B), tertiary (P < 0.001, Fig. 2C), and quaternary (P < 0.01, Fig. 2D) segments of the BLA dendritic neurons compared to the non-stressed animals. Met administration blunted the stress-induced increase in dendritic spine density in the BLA with no effects in any dendritic segments of non-stressed animals (Fig. 2). These results suggest a coordinated role of stress-induced systemic CORT elevation at the time of stress exposure in the late development of anxiety-like behavior and BLA neuroplasticity.

**Figure 2.**
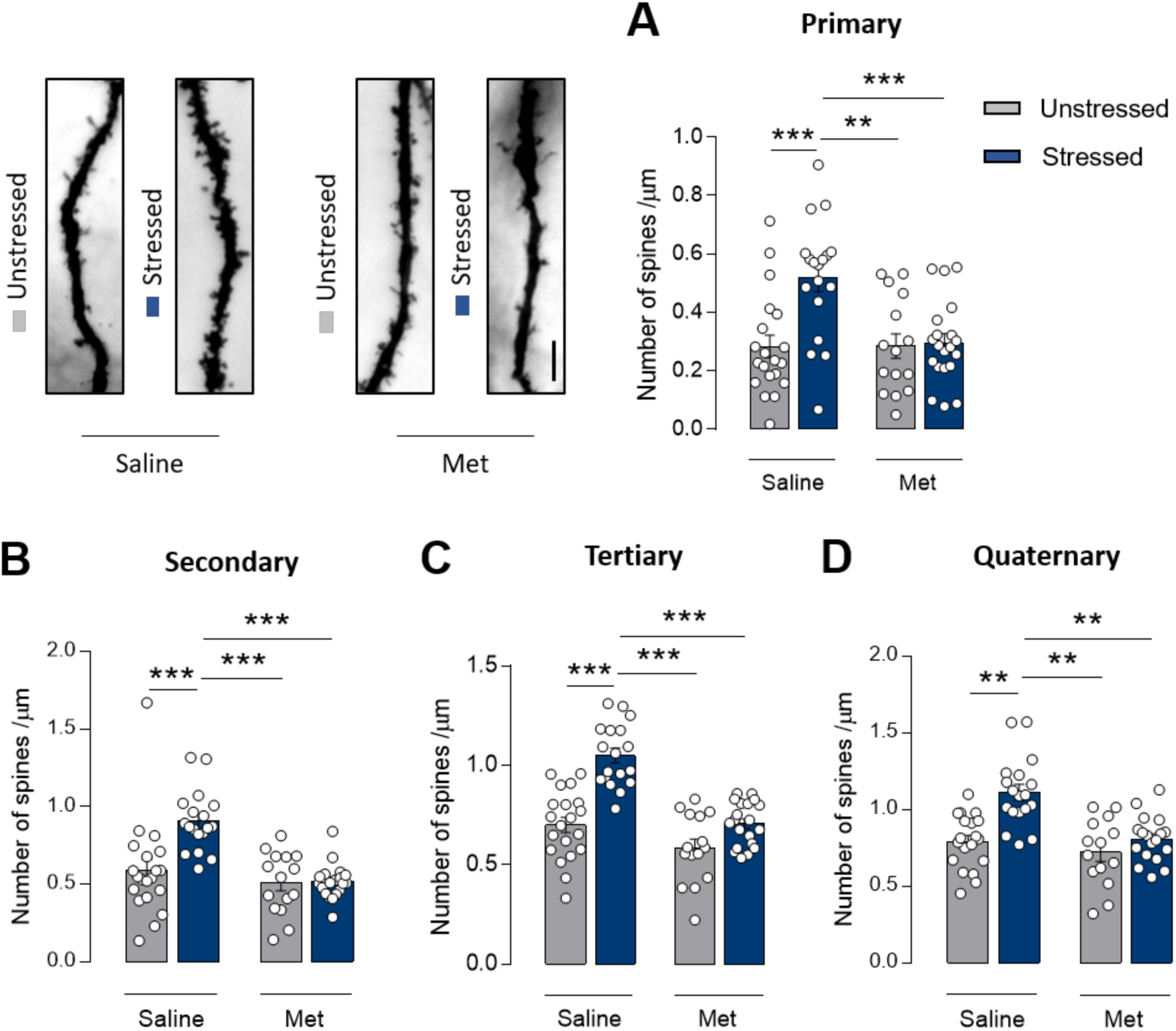
Pre-stress Met administration prevented the stress-induced increase in BLA dendritic spine density. (A-D) Stressed saline-treated animals showed an increase in spine density, an effect prevented by pre-stress Met administration in the primary (A; n = 15–20 neurons for each group, two-way ANOVA: stress F1, 69 = 9.351, P = 0.0032; Met F1, 69 = 7.786, P = 0.0079; stress-Met interaction F1, 69 = 7.72, P = 0.0070), secondary (B; n = 15–20 neurons for each group, two-way ANOVA: stress F1, 69 = 9.871, P = 0.0025; Met F1, 69 = 20.65, P < 0.0001; stress-Met interaction F1, 69 = 8.895, P = 0.0038), tertiary (C; n = 15–20 neurons for each group, two-way ANOVA: stress F1, 69 = 42.17, P < 0.0001; Met F1, 69 = 39.75, P < 0.0001; stress-Met interaction F1, 69 = 9.809, P = 0.0025), and quaternary (D; n = 14–20 neurons for each group, two-way ANOVA: stress F1, 65 = 18.62, P < 0.0001; Met F1, 65 = 15.92, P = 0.0002; stress-Met interaction F1, 65 = 6.267, P = 0.0148) dendrite segments compared to non-stressed animals. (Top, left) Representative photomicrography of a BLA’s secondary dendrite segment. (Scale bar, 10 µm). Results are represented as the mean ± SEM. Tukey’s multiple comparisons post-hoc test). Significant differences between groups are indicated as ** P < 0.01 and *** P < 0.001.

### The delayed effects of acute stress on anxiety-like behavior and BLA dendritic spine density increase depend on neuronal GR genomic activity

Next, we examined whether neuronal GR activation in the BLA was implicated in the late stress outcomes associated with acute stress-induced plasma CORT elevation. Two hours of restraint stress increased GR nuclear translocation into BLA neurons immediately after stress (Fig. S1A and B). Moreover, Western blot analyses revealed an increase in GR nuclear translocation in the BLA (P = 0.0346; Fig. S1C) and phosphorylation at the Ser211 site (P = 0.0166; Fig. S1D), which is associated with a gain in GR transcriptional function (16, 17). Ten days after stress and immediately after the EPM test, there was no difference in the GR nuclear translocation in the BLA of the stressed animals (Fig. S2). Likewise, we found no changes in serum CORT levels of animals submitted to EPM previously stressed ten days before, a result consistent with a previous study (8).

These findings collectively suggest that CORT signaling through GR at the time of stress ultimately triggers delayed BLA alterations. In line with this, animals infused with the GR antagonist RU486 directly into the BLA immediately before stress (Fig. S3A) displayed a higher percentage of entries (P = 0.0161, Fig. S3B) and time spent (P = 0.0215; Fig. S3C) in the EPM open arms ten days later, with no changes in the total distance traveled (Fig. S3D), compared to stressed animals that received vehicle infusions.

Previous studies reported that a rise in CORT serum levels leads, via GR, to rapid (non-genomic) and long-lasting (genomic) modifications in BLA neuronal excitability and synaptic plasticity (18, 19). Because of the delayed effect of stress-associated GR signaling on anxiety-like behavior and remodeling of dendritic spines in the BLA, we expressed a transcriptionally inactive, truncated form of GR in the BLA, the so-called dominant negative GR (dnGR). Cultured cells infected with dnGR responded to CORT treatment with the receptor’s translocation to the cell nucleus as control GFP-infected cells (Fig. S4A and B). However, even with a high GR expression in the nucleus, dnGR-infected cells expressed fewer GR-related mRNA transcription products than GFP-infected cells (Fig. S4C).

The HSV-based plasmid has a transgene expression peak three to four days after infection, followed by a rapid decline in transgene expression (20). Thus, we exposed the animals to restraint stress four to five days after intra-cranial delivery of the virus (Fig. 3A) to maximize any inhibition of genomic GR activity during the stress session. The behavioral test was administered ten days later, a time point at which viral expression of the transgene should be minimal.

**Figure 3.**
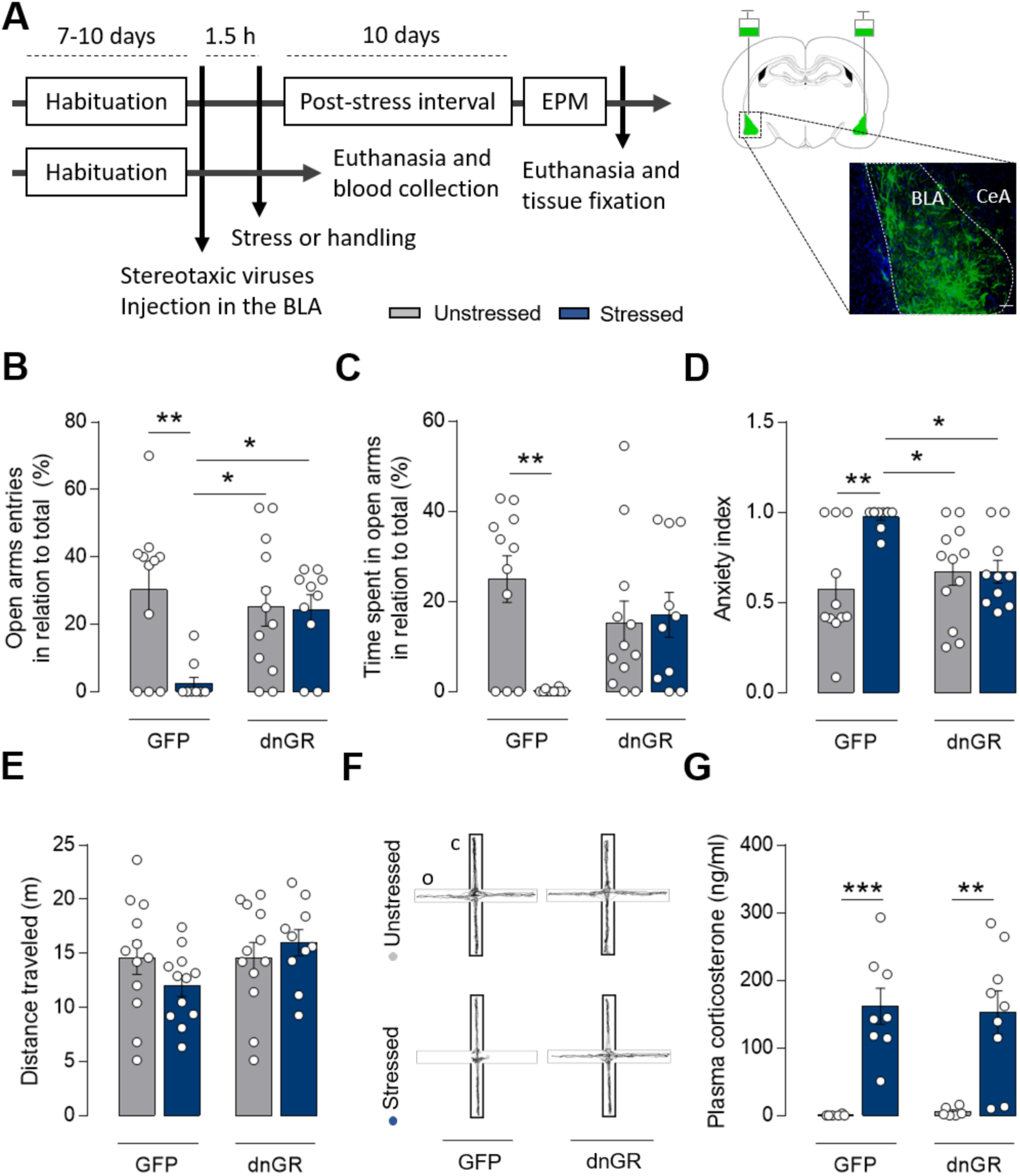
dnGR overexpression in the BLA prevented the delayed emergence of stress-induced anxiety-like behavior. (A, left) Schematic representation of the experimental design shows the virus injection timeline into BLA (GFP or dnGR), the stress submission, and the EPM behavioral test. A cohort of animals was euthanized immediately after the end of the stress session. (Right) Representative figure showing the viral injection into BLA and photomicrography of an infected BLA (GFP, green). (Scale bar, 100 µm). CeA: central nucleus of the amygdala. (B) GFP-stressed animals exhibited reduced open arm entries in the EPM test compared to GFP-non-stressed and stressed and non-stressed animals overexpressing dnGR in the BLA (n = 10–12 animals for each group, two-way ANOVA: stress F1, 39 = 7.527, P = 0.0091; dnGR F1, 39 = 2.612, P = 0.1114; stress-dnGR interaction F1, 39 = 6.724, P = 0.0133). (C) GFP-stressed animals spent less time in the EPM open arms than GFP-non-stressed animals. dnGR animals (stressed and non-stressed) exhibited no differences in time spent in open arms compared to GFP non-stressed animals (n = 10–12 animals for each group, two-way ANOVA: stress F1, 39 = 6.595, P = 0.0142; dnGR F1, 39 = 0.6368, P = 0.4297; stress-dnGR interaction F1, 39 = 8.864, P = 0.0050). (D) Stress increased the anxiety index in GFP-expressing but not in dnGR-expressing rats (n = 10–12 animals for each group, two-way ANOVA: stress F1, 39 = 7.943, P = 0.0075; dnGR F1, 39 = 2.06, P = 0.1592; stress-dnGR interaction F1, 39 = 8.047, P = 0.0072). (E) Neither stress nor BLA virus infection influenced the animal’s locomotor activity in the EPM (n = 10–12 animals for each group, two-way ANOVA: stress F1, 42 = 0.212, P = 0.6476; dnGR F1, 42 = 2.257, P = 0.1405; stress-dnGR interaction F1, 42 = 2.283, P = 0.1383). (F) Representative tracking plots for the average motion of each experimental group in the EPM. (G) Acute restraint stress increases CORT plasma levels in GFP- and dnGR-overexpressing animals immediately after the stressor stimulus (n = 6–9 animals for each group, two-way ANOVA: stress F1, 25 = 36.25, P < 0.0001; dnGR F1, 25 = 0.007222, P = 0.9330; stress-dnGR interaction F1, 25 = 0,07801, P = 0.7823). Results are represented as the mean ± SEM. Tukey’s multiple comparisons post-hoc test. Significant differences between groups are indicated as * P < 0.05, ** P < 0.01, and *** P < 0.001.

Stressed GFP-infected animals exhibited anxiety-like behavior, as evidenced by reduced entries (P < 0.01; Fig. 3B) and time spent (P < 0.01; Fig. 3C) in open arms compared to the GFP-infected non-stressed animals. On the other hand, dnGR expression in the BLA prevented the anxiogenic effect of acute stress. Even when stressed, these animals displayed similar levels of open arm exploration compared to the non-stressed GFP-infected group and significantly increased open arm exploration relative to the stressed GFP-infected animals. This situation was observed for open arm entries (P < 0.05) and the anxiety index (P < 0.05) (Fig. 3B and D). There were no effects of stress or dnGR expression on distance traveled in the maze, suggesting that differences in motor activity did not contribute to the observed behavioral differences (Fig. 3E and F). Additionally, dnGR expression did not impact the release of CORT during acute stress. Similar stress-induced levels of CORT were observed in the stressed dnGR- and GFP-infected groups (Fig 3G) (P < 0.001 and P < 0.01, respectively). Thus, the blunting of the delayed behavioral consequences of stress by dnGR is not due to altered CORT release.

We measured the GFP protein signal throughout all subcellular compartments in brain sections taken from the animals after the EPM and analyzed spine density to determine whether dnGR expression prevented stress-induced changes in BLA spine density. As shown in Fig. 4, there was a stress-induced increase in the dendritic spine density in the primary (P < 0.05; Fig. 4A), secondary (P < 0.05; Fig. 4B), and tertiary (P < 0.001; Fig. 4C) dendritic segments of GFP-infected animals compared to non-stressed animals. Interestingly, the overexpression of dnGR in the BLA prevented the stress-induced increase in the spine densities in the secondary (P < 0.05; Fig. 4B) and tertiary (P < 0.001; Fig. 4C) dendritic segments but not in the primary ones (Fig. 4A).

**Figure 4.**
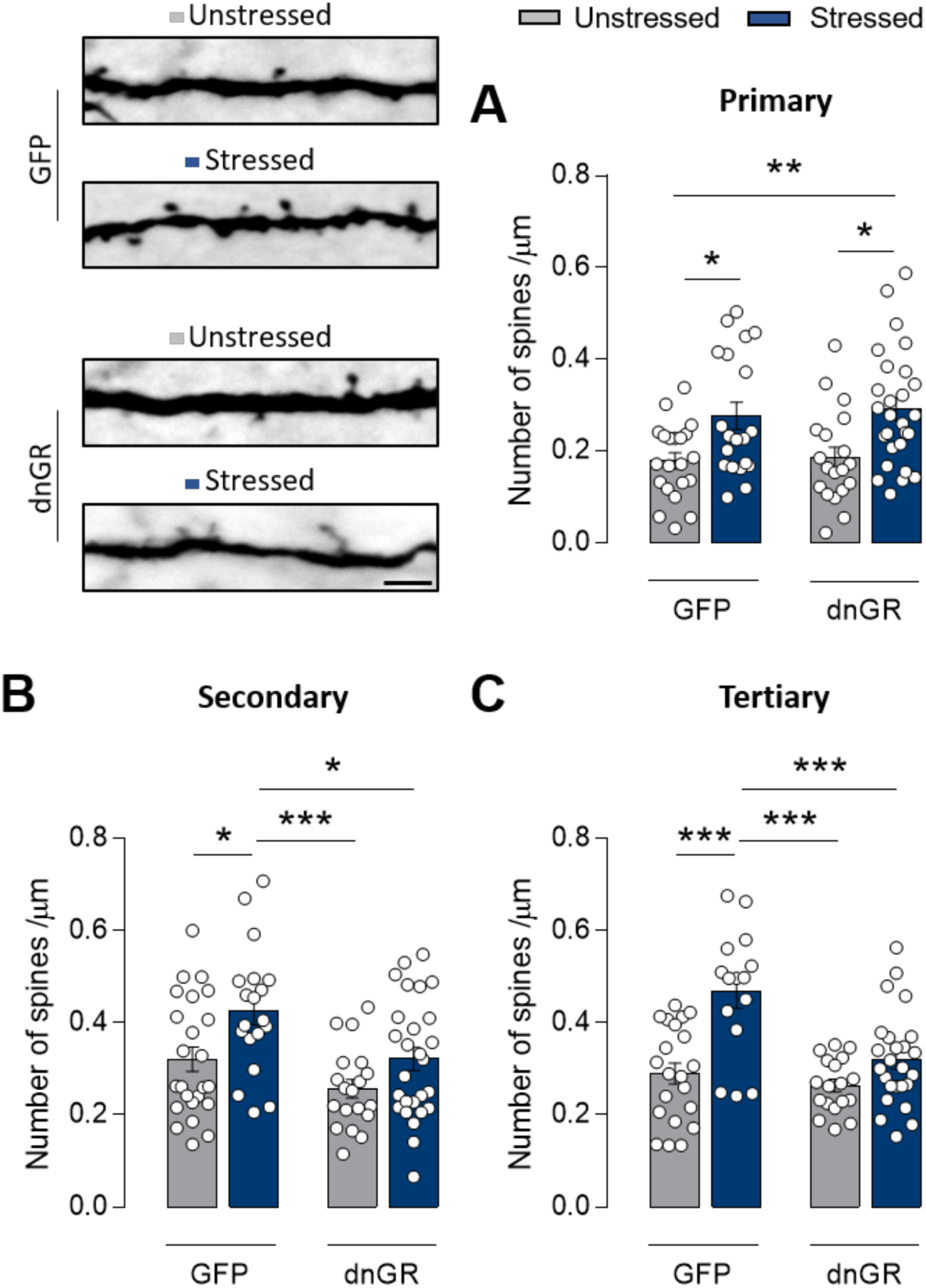
dnGR expression in the BLA prevented the stress-related increase in BLA’s dendritic spine density. (A-C) GFP-stressed animals showed an increase in spine density, an effect prevented by the dnGR overexpressing in the BLA in the primary (A; n = 19–27 neurons for each group, two-way ANOVA: stress F1, 83 = 17.32, P < 0.0001; dnGR F1, 83 = 0.188, P = 0.6657; stress-dnGR interaction F1, 83 = 0.02408, P = 0.8771), secondary (B; n = 19–27 neurons for each group, two-way ANOVA: stress F1, 85 = 10.49, P = 0.0017; dnGR F1, 85 = 10.34, P = 0.0018; stress-dnGR interaction F1, 85 = 0.6021, P = 0.4399), and tertiary (C; n = 15–25 neurons for each group, two-way ANOVA: stress F1, 69 = 16.8, P = 0.0001; dnGR F1, 69 = 8.421, P = 0.0050; stress-dnGR interaction F1, 69 = 7.85, P = 0.0066) dendrite segments compared to non-stressed animals. (Top, left) Representative photomicrography of a BLA’s secondary dendrite segment. (Scale bar, 10 µm). Results are represented as the mean ± SEM. Tukey’s multiple comparisons post-hoc test. Significant differences between groups are indicated as * P < 0.05, ** P < 0.01, and *** P < 0.001.

### An increase in GR activity in the BLA is sufficient to induce a delayed anxiogenic effect

We stereotaxically injected dexamethasone (Dex) into the BLA. Ten days later, we evaluated the anxiety-related behavior (Fig. 5A) to assess whether acute elevation of CORT in the BLA was sufficient to induce delayed anxiety. The EPM test results showed that Dex infusion attenuated the open-arm entry percentage (P = 0.0482; Fig. 5B) and time spent in the open arms (P = 0.0285; Fig. 5C), and elevated anxiety index (P = 0.0127; Fig. 5D), compared to stressed saline-infused animals. Since the Dex-treated animals moved less in the maze than the saline ones (P = 0.0048; Fig. 5E and F), we performed a second anxiety-related behavioral test, the light/dark box test. Since Dex-treated animals spent less time in the light compartment than saline-administered animals, this test confirmed that Dex-treated animals displayed more remarkable anxiety-like behavior than saline-treated animals (P = 0.0369; Fig. 5G).

**Figure 5.**
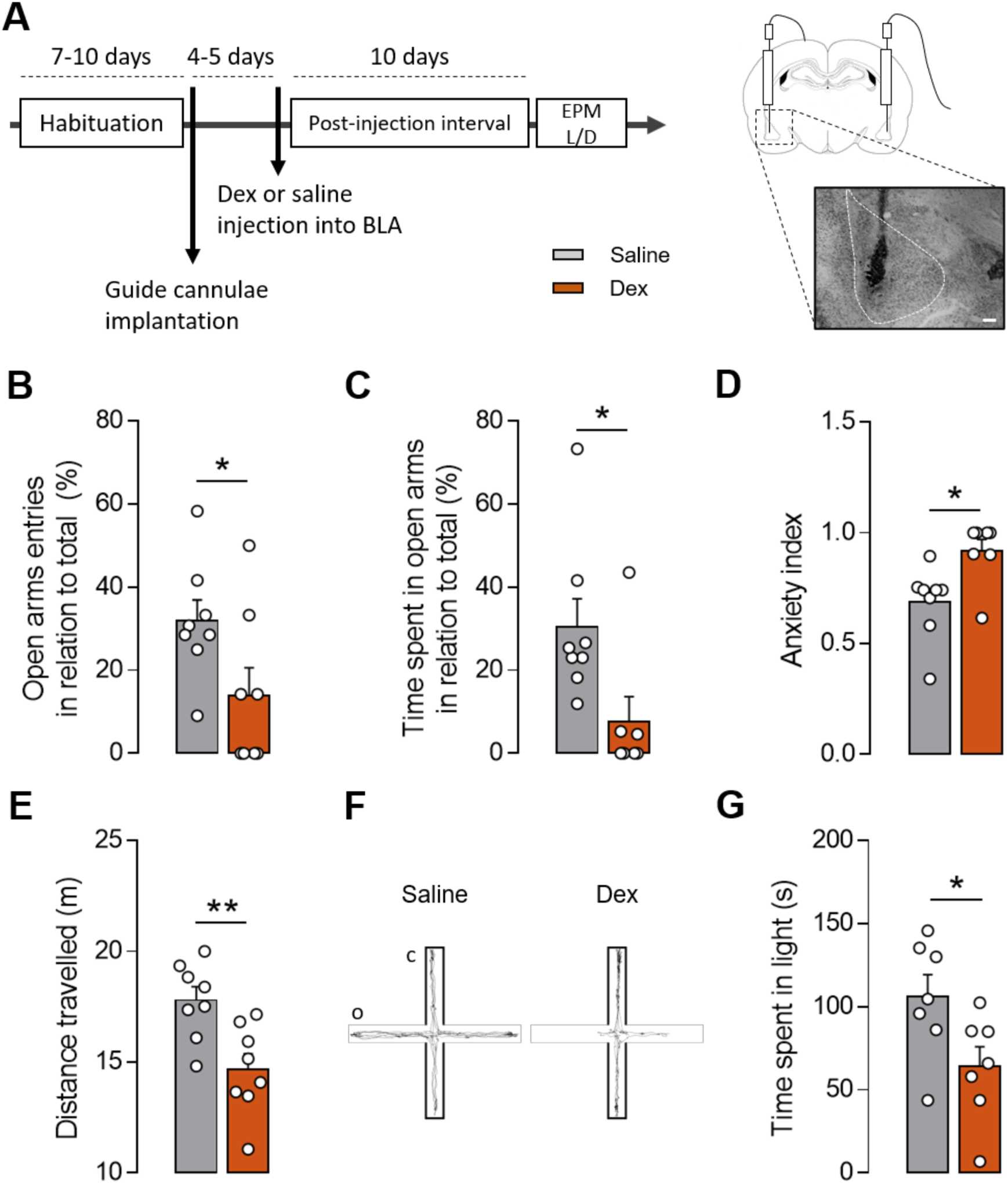
Intra-BLA dexamethasone administration promotes a anxiety-like behavior ten days after infusion. (A, left) Schematic representation of the experimental design showing the timeline of dexamethasone (Dex) or saline injection into BLA, the stress submission, and the behavioral tests. (Right) Representative figure and photomicrography showing a representative location of cannulae and injection needle tips into BLA. (Scale bar, 100 µm); (B–D) Intra-BLA dexamethasone administered animals exhibited reduced open arms entries (B; n = 8 animals per group, two-tailed student’s t-test: P = 0.0482) and time spent (C; n = 7- 8 animals per group, two-tailed student’s t-test: P = 0.0285), as well as increased anxiety index (D; n = 7-8 animals per group, two-tailed student’s t-test: P = 0.0127) in the EPM test comparing to saline administered animals. (E) Intra-BLA dexamethasone promoted a delayed reduced motor activity in the EPM test (n = 8 animals per group, two-tailed student’s t-test: P = 0.0048). (F) Intra-BLA dexamethasone administration promoted a delayed decrease in time spent in the lit compartment comparing to saline administered animals in the light/dark (L/D) test (n = 7 animals per group, two-tailed student’s t-test: P = 0.0369). (G) Representative tracking plots for the average motion of each experimental group in the EPM. Results are represented as mean ± SEM. Significance differences between groups are indicated as * P < 0.05.

### BLA beta-adrenergic receptor activation is necessary for the expression of delayed stress-related anxiety-like behavior

The acute anxiogenic effect of NE is widely known and involves the noradrenergic projections from the locus coeruleus (LC) to the BLA (21). However, whether NE contributes to delayed anxiogenic effects is not known. We addressed this question by infusing the alpha-2 adrenergic receptor antagonist yohimbine into the BLA and examining the acute and delayed effects on anxiety-like behavior (Fig. 6A). Previous studies reported that antagonizing alpha-2 autoreceptors in the BLA increased firing from noradrenergic nerve terminals (12, 13). Consistent with previous reports, we observed that increased NE activity in the BLA produced an acute anxiogenic effect in the open field test (Fig 6B, C, D, E). Additionally, intra-BLA administration of yohimbine did not induce anxiety-like behavior in the EPM test ten days later (Fig. 6F, G, H, I).

**Figure 6.**
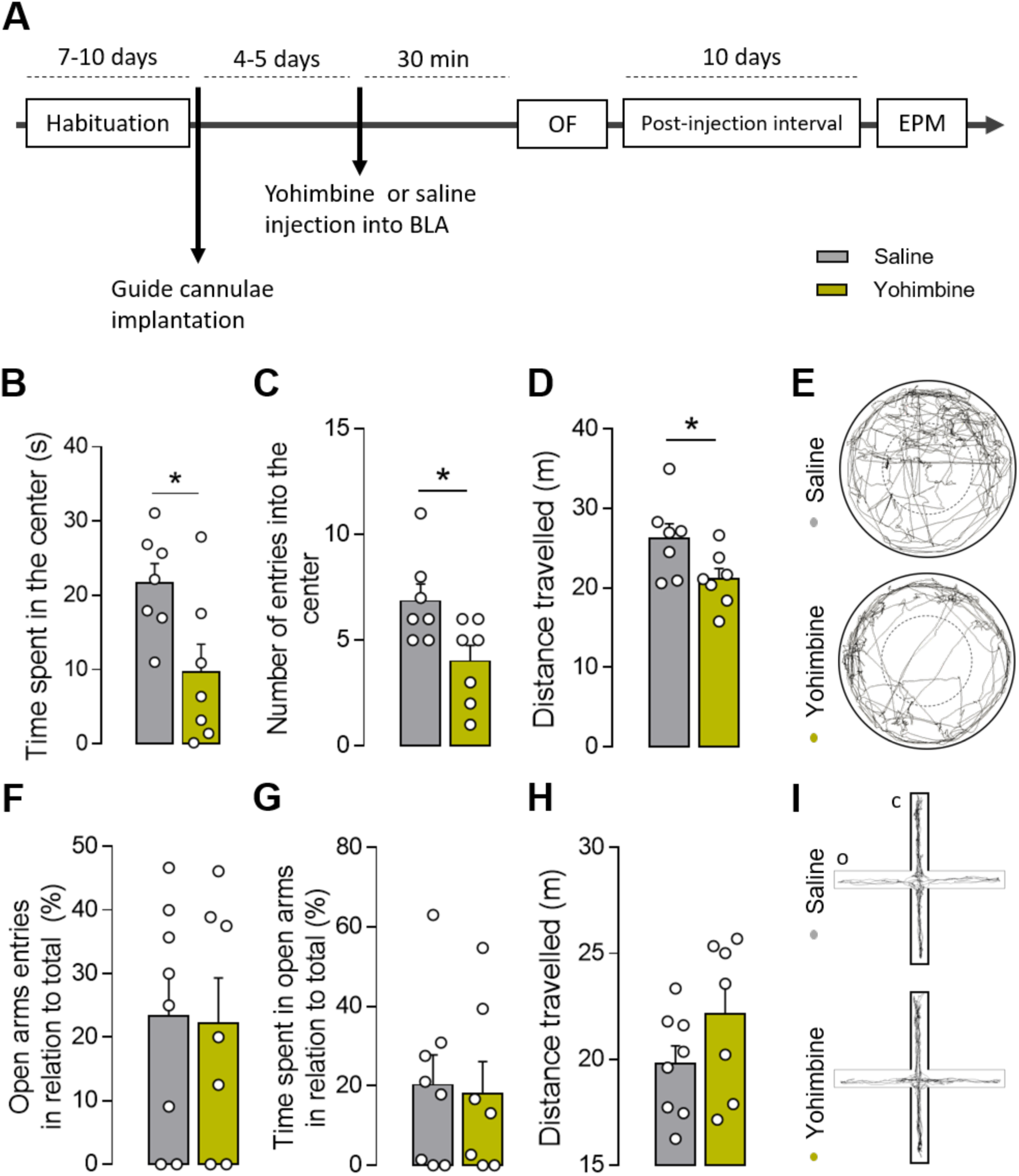
Intra-BLA yohimbine administration promotes an acute, but not delayed, anxiety-like behavior. (A, left) Schematic representation of the experimental design showing the timeline of yohimbine or saline injection into BLA and the behavioral tests. (Right) Representative figure and photomicrography showing a representative location of cannulae and injection needle tips in the BLA. (B-C) Animals that received intra-BLA yohimbine exhibited reduced time spent (B; n = 7 animals per group, two-tailed student’s t-test: P = 0.0225) and entries (C; n = 7 animals per group, two-tailed student’s t-test: P = 0.0234) in the center area of the OF. (D) Intra-BLA yohimbine promoted an acute decrease in motor activity in the OF (n = 7 animals per group, two-tailed student’s t-test: P = 0.00462). (E) Representative tracking plots for the average motion of each experimental group in the OF. (F-G) There were no differences in the open arms entries (F; n = 7-8 animals per group, two-tailed student’s t-test: P = 0.9058) and time spent (G; n = 7-8 animals per group, two-tailed student’s t-test: P = 0.8491) in the EPM test of BLA yohimbine administered animals comparing to saline administered animals. (H) Intra-BLA yohimbine did not influence the animal’s locomotor activity in the EPM (n = 7-8 animals per group, two-tailed student’s t-test: P = 0.1617). (I) Representative tracking plots for the average motion of each experimental group in the EPM. Results are represented as mean ± SEM. Significance differences between groups are indicated as * P < 0.05.

Because increasing NE activity in the BLA has an acute, but not delayed, anxiogenic effect, we examined whether beta-adrenergic receptor activity is critical for the delayed manifestation of anxiety-like behavior in previously stressed animals. Thus, we administered saline or propranolol, a beta-adrenoceptor antagonist, into the BLA of the animals before the behavioral test, ten days after stress (Fig. 7A). As shown in Fig. 7, stressed propranolol-treated animals presented a higher percentage of open arm entries (P < 0.05; Fig. 7B) and time spent in the EPM (P < 0.05; Fig. 7C) than stressed saline-administered animals. These effects were not accompanied by changes in the animals’ locomotor activity (Fig. 7D). Therefore, these results suggest that even at a delayed time point at which stress-dependent increases in spine density are present (Fig. 2), stress-related changes in anxiety still depend on beta-receptor activation in the BLA during the behavioral task.

**Figure 7.**
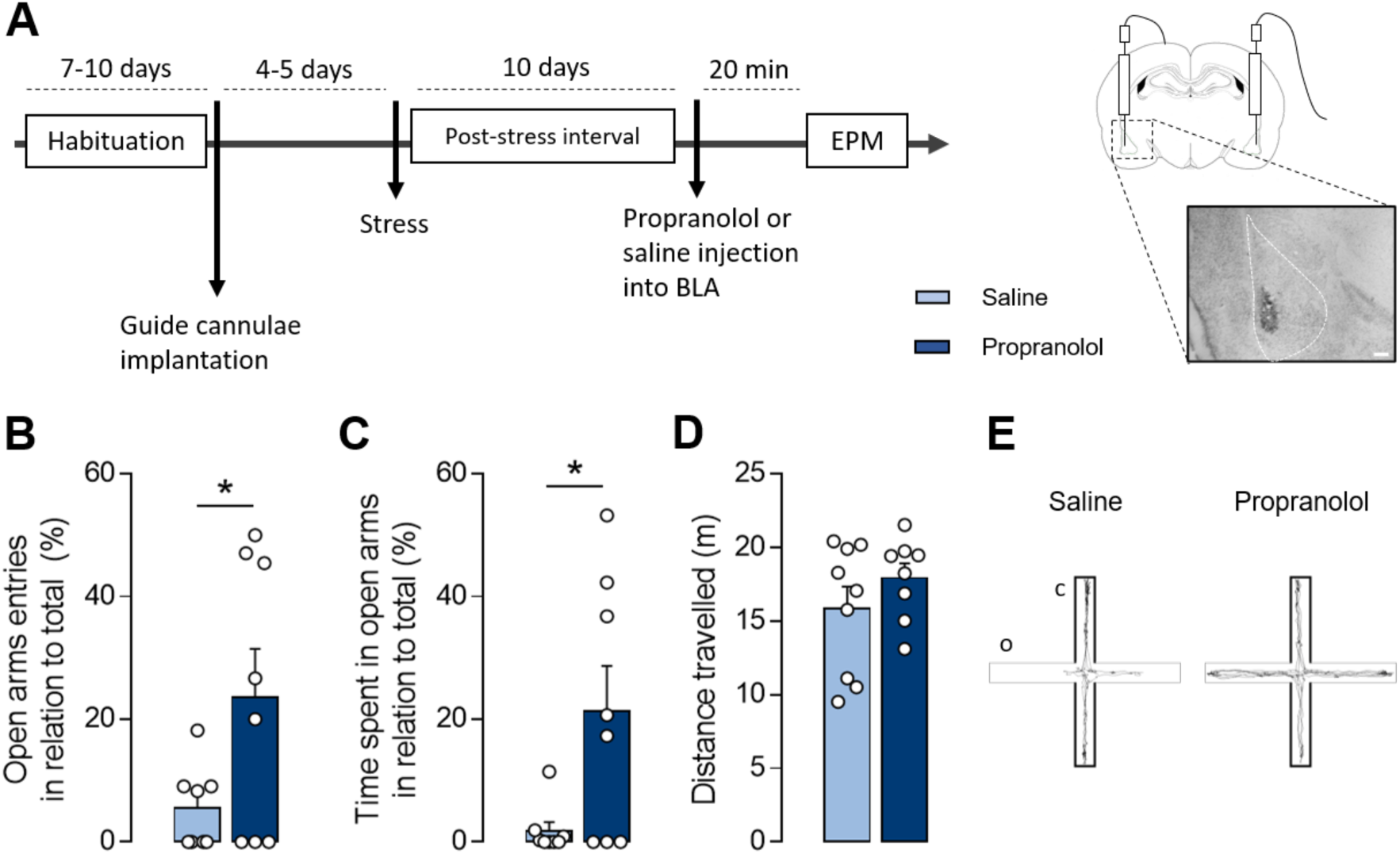
Pre-EPM test propranolol administration in the BLA prevented the delayed manifestation of the stress-induced anxiety-like behavior. (A, left) Schematic representation of the experimental design shows the propranolol or saline injection timeline into BLA, the stress submission, and the EPM test. (Right) Representative figure and photomicrograph showing the representative location of cannulae and injection needle tips into the BLA. (B-E) Stressed, intra-BLA saline-administered animals exhibited reduced open arms entries (B; n = 8 animals per group, two-tailed Student’s t-test: P = 0.0441) and time spent (C; n = 8 animals per group, two-tailed Student’s t-test: P = 0.0218) in the EPM test compared to stressed, intra-BLA propranolol-administered animals. (D) There was no influence of intra-BLA propranolol administration on the animal’s locomotor activity in the EPM (n = 8-9 animals per group, two-tailed Student’s t-test: P = 0.0285). (E) Representative tracking plots for the average motion of each experimental group in the EPM. Results are represented as the mean ± SEM. Significant differences between groups are indicated as * P < 0.05.

## Discussion

Here, we report that inhibiting CORT synthesis during acute stress prevented stress-related behavioral and morphological effects that emerged ten days later. This finding complements a previous study reporting that single systemic administration of CORT, simulating an acute stress exposure, induced a delayed anxiogenic effect, along with parallel increases in dendritic length, branch points, and spine density in BLA neurons (12, 13). It is important to note that we showed that inhibiting CORT alone (Fig. 1-2), but not catecholamines (Fig. 7, 6), sufficiently hindered these stress effects. In this sense, CORT release is essential for the late effects of stress on anxiety-like behavior and dendritic remodeling of the BLA, a result consistent with others (12, 22).

Amygdala hyperactivity is considered a biological substrate of stress- and anxiety-related disorders in human and animal models (23). Stimulation of BLA projection neurons triggers anxiety-like behavior in rodents (24), while acute and chronic stress increases the intrinsic excitability of BLA pyramidal neurons (25, 26). Indeed, symptoms of anxiety manifested by patients suffering from stress-related disorders, such as PTSD, are correlated with amygdala hyperactivity (27, 28). The greater concentration of synaptic contacts in the dendritic spines, compared to the rest of the neuronal surface, confers heightened ionic conductance in this area, resulting in higher excitatory postsynaptic potentials (EPSPs) amplitude (see (29). It is plausible that the delayed morphological changes in the BLA dendritic tree following acute restraint stress produce a more excitable neuronal phenotype, although this remains untested.

Accordingly, rats exposed to a single prolonged session of stress present the delayed anxiogenic effect, which co-occurs with BLA dendritic tree hypertrophy and increased miniature excitatory and inhibitory postsynaptic frequency currents in this brain area (30). Interestingly, the lack of genomic GR signaling in the BLA neurons prevented the stress-related increase in the spine density in the most distal dendritic branches (Fig. 4B, C) without affecting the most proximal ones (Fig. 4A). This effect is consistent with other reports showing that distal synaptic inputs on BLA neurons are more likely to be excitatory when compared to proximal synapses, probably due to a higher GABAergic input in the latter (31).

A previous study reported that acute elevations in blood CORT levels modulate dendritic spine turnover in the cortex, producing a short-term effect on eliminating and forming new dendritic spines (32). Interestingly, acute stress and a single systematic injection of CORT promote delayed (at least ten days later) but not short-term (one day later) changes in the BLA dendritic tree (7, 22).

Protein synthesis is a critical step for forming new dendritic spines (33). Our findings show that the stress-related anxiety-like behavior and the increase in the BLA dendritic spine density are governed by the direct action of CORT in this brain region, phenomena dependent on genomic GR activity. Notably, the GR mobilization by Dex injection into BLA generated a delayed anxiety-like behavior even without stress. However, the mechanism by which GR influences the origin and stabilization of new spines is not yet fully understood. Nevertheless, growing evidence has shown a synergistic and interdependent action between GR and the TRKB-BDNF receptor in the motor cortex during motor-related learning (34-36).

The role of NE in modulating behavior is well known and is closely related to the immediate effects of changing/increasing the perception of threats during stress exposure (37). Acute stress leads to a rapid release of NE in the limbic system, including the BLA and hippocampus, primarily via projections of noradrenergic neurons from the LC (21, 38). The activation of beta-adrenergic receptors in the BLA promotes increased excitatory synaptic transmission and increased neuronal spike firing rates [see (15)]. In the present study, we confirmed an acute anxiogenic effect of increased availability of NE in the BLA (via yohimbine administration) (Fig. 6). Still, this increase in NE release did not promote delayed behavioral changes. In contrast, Dex administration in the BLA was sufficient for promoting delayed behavioral changes (Fig. 5). Additionally, blocking NE signaling directly into BLA with propranolol at the time of behavioral testing prevented the manifestation of anxious-like behavior in previously stressed animals (Fig. 7).

The synergism of GR and noradrenergic transmission in the BLA has already been critical in potentiating the effects of arousal on memory retention (39). Here, we reported a non-simultaneous but coordinated and (possibly) non-direct dependence of the delayed manifestation of stress-induced anxiety on both GR and beta-adrenoceptor signaling in the BLA. It is important to note that our previous study did not detect increased CORT release ten days after the stress or immediately after the EPM test (8), indicating that elevated CORT levels were not responsible for delayed anxiety-like behavior. We propose that the stress-related delayed plastic changes in the BLA’s dendritic arbor rely on genomic GR signaling events, which may potentiate the stress response to NE. Therefore, given the morphological effects of stress, the delayed behavioral outcome depends on NE signaling via beta-adrenoceptors.

## Materials and Methods

### Subjects

Male Wistar rats (60-days old) were obtained from either the Facility for SPF Rat Production at the Institute of Biomedical Sciences - Animal Facility Network at the University of São Paulo (USP) or Harlan Sprague-Dawley Inc (Indianapolis, IN, USA). Neonatal Wistars rats (1 - 4 days old) were used for the primary cortical culture. Upon arrival, the animals were maintained at the vivarium in the USP Pharmacology Department (Unit 1) or Department of Brain and Cognitive Sciences, Massachusetts Institute of Technology (MIT). The animals were pair-housed in standard polypropylene cages (30 × 40 × 18 cm) with ad libitum access to food and water in a temperature controlled (23 ± 2 °C) room with a 12-hour light-dark cycle (light on at 6:00 am). All experiments were conducted under the standards of the Ethics Committee for Animal Use of the Institute of Biomedical Sciences/USP (CEUA - ICB 85/2016) and by the Committee for Animal Care at MIT. All methods were in accordance with the Guidelines of the Brazilian National Council for the Control of Animal Experimentation (CONCEA) and the US National Institutes of Health (NIH) Guide for the Care and Use of Laboratory Animals.

Upon arrival in the vivarium, the animals were pair-housed in standard polypropylene cages (30 × 40 × 18 cm), provided with food and water ad libitum, and maintained in a temperature controlled (23 ± 2 °C) room with a 12-hour light-dark cycle. All experimental procedures were initiated after a habituation period of 7 to 10 days and performed between 0900 and 1200 during the cycle’s light phase.

## Animal experimental procedures

### Acute restraint stress

The animals were placed in a ventilated PVC restraint tube for two hours in a lighted and soundproof room. The behavioral experiments were conducted ten days after the end of the stress session.

### Behavioral tests

The behavioral tests were conducted in a soundproof room with low light (10-14 lux). All tests were videotaped using a digital video camera (Logitech C920, Lausanne, Switzerland) from the top of the apparatuses. After each session, the apparatuses were cleaned with 5% alcohol to remove olfactory cues.

#### Elevated plus maze

Animals were placed in the center of the maze (elevation of 50 cm, 50 cm × 10 cm each arm), facing one of the open arms, and allowed to freely explore the apparatus for 5 min. We scored entries, time spent in each arm, and the total displacement in the maze using Ethovision software (Noldus, the Netherlands). The anxiety index (40, 41) was calculated as follows: 1 - [(time spent in open arms/total time on the maze) + (number of entries to the open arms/Total exploration in the maze)/2].

#### Light/dark box

Animals were placed in the middle of the light chamber (45 × 41 × 21 cm light chamber, 65 lx; 35 × 41 × 21 cm dark chamber, 0 lx) facing the opposite side of the connecting door. Each trial lasted 5 min, and the duration of time spent in the light chamber was used as a parameter to address anxiety-like behavior.

### Stereotactic surgery

Drugs and viral vectors were injected into the BLA through stereotactic surgery, as described by (42), with some modifications. Rats were anesthetized with isoflurane (4% oxygen saturation for induction and 2-3% for maintenance), received a dose of anti-inflammatory for preemptive analgesia (ketoprofen, 5 mg/kg, subcutaneous), and positioned in the stereotaxic apparatus (Kopf Instruments). The skin was disinfected (70% ethanol, 1% iodine, and 2% glycerin), the skin covering the skull was incised, and small holes were drilled in the skull surface to pass cannulae or injection needles. The stainless-steel guide cannulae (23-gauge, World Precision Instruments, Sarasota, FL, USA) were positioned bilaterally 1.0 mm dorsal to the BLA, using the following coordinates: A/P, -2.4; M/L, ± 5.1; D/V, -6.0 mm relative to bregma and the surface of the brain (43). The cannulae were fixed with polyacrylic cement and anchored to the skull with stainless steel screws. Drug infusions were made through injectors (30-gauge stainless steel that extended 1.0 mm beyond the tip of each guide cannula) after the animal’s complete recovery from cannula implantation surgery (4 to 5 days). A polypropylene tube connected the injectors to a Hamilton syringe (5 μl, Hamilton Co., Reno, NV, USA). The infusion rate was 0.1 μl/min with 1 min for diffusion before the injector was removed from the guide cannula. The viral particles were injected with stainless-steel needles coupled to a Hamilton syringe (5 μl, Hamilton Co.) fixed in an infusion pump (Pump 11 Elite, Harvard Apparatus, Holliston, MA, USA) coupled to the stereotaxic apparatus. The injections were placed bilaterally in the BLA using the following coordinates: A/P, -2.4; M/L, ± 5.1; D/V, -7.0 mm. The viruses were infused at a rate of 0.1 μl/min for 20 min (total of 2 μl per injection site, totaling 2 × 106 viral particles). The needles remained in place for ten minutes before being removed. The injections were performed 4-5 days before stress, a critical time to reach the viral expression peak (44). Only animals with bilateral viral infection in the BLA were considered in the behavioral analysis. After surgery, the animals were under observation for three days. If they showed any sign of pain or discomfort, they received a daily dose of anti-inflammatory (ketoprofen, 5 mg/kg, subcutaneous).

### Cannula placement verification, viral infection visualization, and immunofluorescence

Cannula placement verification was achieved by Nissl staining. The expression of GFP indicated viral infection in the BLA. Animals in which the cannulae were not correctly positioned bilaterally in the BLA, i.e., GFP expression was not restricted to the BLA, were excluded from the study. The GFP detection and immunofluorescence assays are detailed below.

### Drug administration

Metyrapone (i.p.; 150 mg in 1 ml of saline; Sigma-Aldrich) was administered 1.5 hours before restraint stress. Bilateral intra-BLA infusion of RU-486 (10ng in 0.5 µl of 5% DMSO; Sigma-Aldrich), dexamethasone (30 ng in 1 µl of 0.5% ethanol), propranolol (2.5 µg in 0.5 µl of saline), or yohimbine (2.5 µg in 0.5 µl of saline), or their corresponding vehicle controls, were performed via stereotaxic surgery.

### Viral vector construct

Viral vectors containing the dnGR receptor gene sequence were constructed as previously described (20, 45, 46). The dnGR gene construct is a chimera composed of the rat GR alpha subunit (GenBank M14053), containing the ligand-binding domain, fused to the human GR beta subunit (GenBank X03348), which includes the DNA-binding site domain. The viral construct was obtained from the MIT viral core facility (McGovern Institute, MIT). The dnGR was packaged in the HSV p1005 vector (short-term expression), a Herpes Simplex Virus 1 (HSV-1) derived amplicon plasmid with an added transcription cassette expressing GFP (separate promoter and poly(A) (44, 47). The empty vector (lacking the dnGR sequence) was used as a control. The IE 4/5 promoter drives the target gene, while a CMV promoter drives the GFP gene. The two transcription cassettes are in a nose-to-tail orientation. Adherent 2-2 cells were transfected with the plasmid of interest (containing dnGR) using the Lipofectamine LTX reagent (Invitrogen). Twenty-four hours after transfection, the cells were super-infected with Helper Virus 5dl1.2, which has a multiplicity of infection of 0.03. The cells and media were harvested 48 hours later when the virus’s cytopathic effect reached 100%. The solutions were frozen and thawed three times, sonicated to release infectious viral particles, and then centrifuged to clear the media of cell debris. The resulting supernatant was passed onto 2-2 cells. This amplification procedure was repeated twice. After the final sonication and centrifugation, the supernatant was purified on a sucrose gradient, viral bands were centrifuged, and viral pellets were resuspended in 10% sucrose in D-PBST. The vectors’ titration was quantified in Vero cells by counting the cells that express the GFP protein. An average titration of approximately 1 × 106 viral particles per µl was achieved.

### Dendritic spine density analysis

For some experiments, the brains were prepared using the Rapid Golgi Kit (FD NeuroTechnologies, Inc., Columbia, MD, USA) according to the manufacturer’s instructions. An immunohistochemistry assay was performed for other experiments to amplify the GFP signal to analyze the dendritic spine density in the BLA neurons previously infected with the HSV vector.

### Golgi staining and spine density analysis/Dendritic spine density analysis

Brains were incubated in an impregnation solution for 14 days, followed by incubation in a cryoprotectant solution for 48 hours. Samples were coronally sliced (120 μm) using a Leica CM3050-S cryostat (Leica Biosystems) and mounted on gelatin-coated slides. Brain sections were dehydrated in ethanol, diaphonized in xylene, and coverslipped. The whole BLA images were captured in a bright field with the slide scanner Zeiss Axio Scan.Z1 (Zeiss), using a 40× objective. A z-stack with a 410 nm step size was created, and the images were analyzed with the ZEN 2 (Zeiss) software at 600× magnification. At least five neurons per animal (four animals per group) were analyzed according to the following criteria: only neurons with the soma wholly localized in the BLA, heavily impregnated, without truncated dendrites. After neuronal soma identification, we classified each dendritic segment as primary (originating from the soma), secondary (originating from the primary branch), tertiary (originating from the secondary branch), or quaternary (originating from the tertiary branch). Regardless of the morphologic classification, all protrusions originating from a dendrite were considered spines. After we determined the origin and the end of each branch, the dendritic branch length was calculated by the software.

### Brain tissue processing for immunofluorescence and immunohistochemistry

Brains were sectioned in a cryostat (Leica CM3050-S, Leica Biosystems, Wetzlar, Germany) in a coronal plane (40-µm-thick) at -22 °C and stored in an anti-freeze solution (0.05 M sodium phosphate buffer, 150 g of sucrose, and 300 ml of ethylene glycol) at -20 °C.

We performed an immunofluorescence assay to visualize GR expression in the BLA. Brain slices were incubated in blocking solution (0.02 M potassium phosphate solution containing 0.3% Triton X-100 and 1% bovine serum albumin) for 1 hour, followed by incubation in blocking solution containing the primary antibody (rabbit anti-GR, dilution 1: 1000, Santa Cruz; mouse anti-MAP-2, dilution 1: 5000, Sigma-Aldrich) for 18 hours (4 °C). The slices were then incubated for two hours (room temperature) in a 0.02 M potassium phosphate solution containing the secondary donkey antibody conjugated to a fluorochrome (AlexaFluor 594 anti-rabbit, dilution 1: 500 or AlexaFluor 488 anti-mouse, dilution 1: 500, Thermo Fisher Scientific). Brain slices were mounted on positively charged slides (Fisherbrand Superfroast Plus, Thermo Fisher Scientific) and covered with a coverslip using Fluoromount-G as the mounting medium (SouthernBiotech, Birmingham, AL, USA). The images were acquired in a confocal microscope (Nikon A1R +, Nikon Instruments Inc., Melville, NY, USA), sectioned in the Z plane at 0.6 µm.

Although the HSV vector carries the gene that encodes GFP, an immunohistochemistry procedure was performed to intensify the signal and improve GFP particle detection throughout the BLA dendritic tree. Brain slices were incubated in blocking solution (0.02 M potassium phosphate solution containing 0.3% Triton X-100 and 3% normal donkey serum) followed by incubation in blocking solution containing the primary antibody anti-GFP raised in rabbit (dilution 1: 1000; Thermo Fisher Scientific) for 18 hours at 4 °C. The slices were then incubated in a 0.02 M potassium phosphate solution containing a biotinylated donkey antibody (dilution 1: 200; Vector) and 0.3% Triton X-100 for two hours at room temperature. The antigen-antibody complex was visualized with the biotin-avidin technique (Elite kit; Vector Laboratories, Burlingame, CA, USA) and 3,3’-diaminobenzidine (DAB) as a chromogen through a peroxidase reaction composed of a 2.5% solution of nickel sulfate and 0.003% hydrogen peroxide in 0.02 M potassium phosphate buffer. The sections were mounted on gelatin-coated slides, air-dried, and dipped in a 0.05% aqueous solution of osmium tetroxide for 20 seconds to enhance the labeling’s visibility, dehydrated, transferred into xylene, and coverslipped with DPX. Spine density was quantified under a bright field, as detailed for the Golgi staining assay.

### Western Blot

GR expression and its phosphorylated form were analyzed by western blot as previously (41).

#### Protein extraction

Cytosolic and nuclear protein extracts were obtained using the Cel-LyticTM NuCLEARTM Extraction Kit (Sigma-Aldrich). BLA tissues stored at -80 °C were thawed and mechanically homogenized with a conical plastic pestle (Thermo Fisher Scientific) in cold lysis buffer (10 mM HEPES, pH 7.9, 10 mM KCl, 1.5 mM MgCl2, 0.5 mm PMSF, 1 mm DTT) supplemented with protease and phosphatase inhibitors (Halt Protease Inhibitor Cocktail, ThermoFisher Scientific), and centrifuged for 20 min at 11,000 × g at 4 °C. The supernatant (cytosolic fraction) was collected in a tube and stored at -80 °C. The remaining pellets were homogenized in extraction buffer (20 mM HEPES, pH 7.9, 300 mM NaCl, 1.5 mM MgCl2, 0.25 mM EDTA, 0.5 mM PMSF, 0.5 mM DTT 25% glycerol) supplemented with protease and phosphatase inhibitors, kept on ice for 30 min, and centrifuged for 5 min at 20,000 × g at 4°C. The resulting supernatants (nuclear proteins) were stored at -80 °C. Protein concentration was determined using the Bradford method (Bio-Rad Laboratories, Hercules, CA, USA).

#### Electrophoresis

Protein samples (20 μg/lane) were size-separated in 10% SDS-PAGE gel (90 V) and then blotted onto Immobilon® PVDF membrane (EMD Millipore Corporation). The immunoblots were stained Ponceau to ensure equal protein loading (48). Blots were blocked with 5% BSA diluted in TBS-T buffer (50 mM Tris-HCl, 150 mM NaCl, 0.1% Tween 20, pH 7.5) for one hour at room temperature and subsequently incubated overnight at 4 °C with specific primary antibodies: monoclonal rabbit anti-GR (dilution 1:1000, Santa Cruz) or polyclonal rabbit anti-Ser211phospho-GR (dilution 1:1000, Cell Signaling). After incubation with the primary antibodies, membranes were probed with a secondary antibody conjugated to horseradish peroxidase (dilution of 1:2000, Kirkegaard & Perry Laboratories, Gaithersburg, MD, USA) for two hours at room temperature. The blots were developed using the Luminata Forte Western HRP substrate (Merck Millipore, Darmstadt, Germany), and the signals were recorded with the ChemiDoc system (Bio-Rad Laboratories). Several exposure times were performed to ensure the linearity of the band intensities. Each band’s relative density was normalized to the value of α-Tubulin (sc-5286, dilution: 1:20,000, Santa Cruz Biotechnology, Dallas, TX, USA).

### Primary cortical cell culture

Neonatal rats were sacrificed by decapitation 1-4 days post-natal. The frontal cortex was dissected in sterile medium (Hank’s buffer: 160 mM NaCl, 5.3 mM KCl, 0.44 KH2PO4, 0.33 mM Na2HPO4, 5.5 mM glucose, 4.0 mM NaHCO3) supplemented with antibiotics (penicillin/streptomycin, Invitrogen), at pH 7.4. The isolated frontal cortex was maintained in Hank’s buffer containing 0.05% trypsin (Vitrocell, Campinas, SP, Brazil) and 0.001% DNAse (Sigma-Aldrich) at 37 °C for 10 minutes. Then the samples were then transferred to Dulbecco’s Modified Eagle Medium culture medium (DMEN; Vitrocell) plus 10% fetal bovine serum (FBS, Vitrocell) and 2 mM glutamine (Sigma-Aldrich), in which it was gently mechanically grounded with the aid of a Pasteur pipette. The solution was filtered through a 70 µM membrane (Becton-Dickinson), and the cells were further cultured in a 24-well plate and incubated at 37 °C with 5% CO2 in a humidified atmosphere (80% relative humidity). Cells were cultured on borosilicate coverslips covered with poly-L-lysine (0.01%, Sigma-Aldrich) in a 24-well plate. For the immunofluorescence assays, the cells were counted before plating using the Trypan Blue exclusion method (Sigma-Aldrich) to calculate the number of viral particles necessary for efficient infection. The cells were maintained in DMEN supplemented with antibiotics and 10% FBS. The medium was changed on the second and fifth days after culturing and maintained until the tenth day.

### Infection of cells by viral particles and analysis of sub-localization and genomic activity of GR

Culture cells were infected with the viral particles (HSV-dnGR and HSV-GFP, MOI of 1) diluted in culture medium (DMEN + 2% FBS) on the eighth culture day to check the sublocalization of the dnGR receptor in the cell compartments in response to CORT. Forty-eight hours later (i.e., the tenth day of culture), the cells were treated with 10 µM CORT in culture medium (DMEN + 2% FBS) for one hour. Then, the medium was removed, and the coverslips were washed with PBS and fixed with 4% formaldehyde (Amresco) for 20 minutes at room temperature. After five washes with PBS, the cells adhered to the coverslips were incubated in a blocking solution (5% donkey serum, 0.05% Triton X-100 in PBS) for two hours at room temperature. Next, the cells were incubated in a blocking solution containing the primary anti-GR antibody (1: 1000, Santa Cruz) for 24 hours and then, after washing with PBS, incubated in PBS containing the secondary antibody (AlexaFluor 594, 1: 2000) and 0.05% Triton X-100 for two hours. Finally, the coverslips were incubated in DAPI solution (1: 100,000, Sigma-Aldrich; PBS with 0.05% Triton X-100) for 20 minutes, washed with PBS, and mounted on slides in Fluoromount-G mounting medium (SouthernBiotech). The slides were analyzed under a fluorescence microscope (Nikon Eclipse 80i; Nikon Instruments Inc).

The polymerase chain reaction (PCR) assay was performed to amplify the Gilz and Fkbp5 genes described below and analyze the genomic GR activity in dnGR-infected cells. The cells were obtained using the same protocol for image analysis, but the cells were not cultured on coverslips. Instead, at the end of the treatment with CORT, the cells were washed five times with PBS and collected in TRIzol® (0.75 µl per sample; Thermo Fisher Scientific). RNA extraction was performed following the manufacturer’s instructions.

### Polymerase chain reaction (PCR)

After solubilizing the cells in TRIzol®, 175 µl of chloroform were added to each sample, followed by incubation at room temperature for three minutes. The samples were centrifuged at 12000 × g for 15 minutes at 4 °C, forming three phases. The first phase contained the total RNA and was transferred to another tube. Next, 5 µg of RNase-free glycogen and 0.5 ml of isopropanol alcohol were added for RNA precipitation. The samples were incubated at room temperature for 10 minutes and centrifuged at 12000 × g for 10 minutes at 4 °C. After discarding the supernatant, 1 ml of 75% ethanol was added to the resulting pellet, followed by vigorous vortexing and centrifugation at 7500 × g for 5 minutes at 4 °C. After discarding the supernatant, the pellet was dried at room temperature, diluted in DEPC water (20 µl), and stored at -80 °C. Two µl of the RNA was used for spectrophotometric quantification at 260 and 280 nm in a Synergy HT Multi-Mode Microplate Reader (BioTek Instruments, Inc, Winooski, VT, USA).

The cDNA was obtained using the SuperScript III Reverse Transcriptase Kit (Thermo Fisher Scientific) according to the manufacturer’s instructions. For the reaction, 12 µl of final solution were used, containing 5 µg of RNA, 1 µl of dNTPs (0.5 nM each), 1 µl of Random Primer in sterile water. The samples were incubated for 5 minutes at 70 °C, and then 8 µl of the Mix solution was added to the solution, resulting in a final volume of 20 µl. Samples were incubated for 10 minutes at 25 °C, 42 °C for 70 minutes, and 70 °C for 15 minutes to denature the enzyme.

For the amplification, 2 µl of the cDNA (20 ng / µl), 5 µl of Amp Buffer, 0.75 ul MgCl2, 0.5 µl of dNTPs, 1.5 µl of the forward primer, 1.5 µl of the reverse primer, 0.5 µl of Taq Polymerase (Promega Co), and Milli-Q water (Merck Millipore) sufficient for 25 µl were added to a microtube. The reaction conditions in the thermal cycler (LifeECO, Bulldog Bio, Inc) were: 95 °C for 2 initial minutes, followed by 30 cycles at 95 °C for 1 minute, 55 °C for 30 seconds, and 72 °C for 1 minute. After the last cycle, the samples remained at 72 °C for 5 minutes. The product was separated using agarose gel electrophoresis (2%), stained with BlueGreen (Thermo Fisher Scientific), and visualized with a ChemiDoc Imager equipped with a UV transilluminator (Bio-Rad Laboratories, Inc). The sequences of the primers used are as follows:

Gapdh: forward, 5’-GGTGTGAACGGATTTGGC-3’ reverse, 5’-CTGG-AAGATGGTGATGGGTT-3’
Gilz: forward, 5’-TGGGGCCTAGTAACACCAAG-3’ reverse, 5’-GAGCACACTGGCATCACATC-3’
Fkbp5: forward, 5’-GAACCCAATGCTGAGCTTATG-3’ reverse, 5’- ATGTACTTGCCTCCCTTGAAG-3’

## Euthanasia of the animals and obtaining the analyzed material

### Guillotine beheading, BLA dissection and blood collection

The animals were deeply anesthetized with isoflurane (maximum exposure of 30 s) and euthanatized by decapitation. The brain was quickly removed in a cold solution of saline-phosphate buffer (137 mM NaCl, 2.68 mM KCl, 1.27 mM KH2PO4, 8.06 mM Na2HPO4). The BLA was bilaterally dissected on a cold glass plate and frozen in a conical tube (1.5 ml) on dry ice. The brain structures were stored at –80 °C for later use. Brains were placed in a 1 mm thick coronal cut acrylic brain matrix (Zivic Laboratories Inc., New Castle, PA, USA), and the dissection was performed under a stereomicroscope (Tecnival – Prolab, Diadema, SP, Brazil). Blood was obtained from the animals’ trunks at the time of decapitation.

### Transcardiac perfusion

The animals were deeply anesthetized with a solution of ketamine hydrochloride (9 mg / 100 g, i.p.) and xylazine (1 mg / 100g, i.p.) and infused transcardiacally using a peristaltic pump (Cole Parmer, Vernon Hills, IL, USA), with 200 ml of 0.9% saline followed by about 500 ml of 4% formaldehyde solution in sodium phosphate buffer (pH 7.4) at 4 °C for 20 minutes. After three hours, the brain was removed from the skull and kept in a 30% sucrose solution in potassium phosphate buffer (0.02 M) at 4 °C for 48 hours. After this cryoprotection period, the brain was removed from the sucrose solution and frozen at -80 °C until histological processing.

### Blood collection and plasma obtaining

Blood was sampled from the body trunk after decapitation, immediately after stress termination. Plasma was obtained by adding EDTA (4.5 mM) to the collected blood and centrifugation at 4000 × g for 10 minutes. Samples were stored at -80 °C.

## Analysis of the concentration of corticosterone in the blood

The Corticosterone EIA kit (Enzo Life Sciences, Inc, Farmingdale, NY, USA) was used to analyze blood CORT levels, following the manufacturer’s guidelines. Plasma samples were diluted at 1:30 in the assay buffer provided by the kit to fit the standard curve’s optimal range. The values depicted in the figures were corrected for the dilution factor.

## Statistics

Statistical analysis was performed using two-way ANOVA. Post hoc Tukey’s multiple comparison test examined differences between individual groups. An unpaired Student’s t-test was used to compare two experimental groups when appropriate. Sample size and animal numbers were estimated based on previous studies, and the ROUT method checked the outlier samples. Quantification of data was blind for all experiments. GraphPad Prism software 9.0 was used for the statistical analysis, and the level of statistical significance was p < 0.05. Data are presented as the mean ± SEM. Detailed statistical information for each test performed is reported in the figure legends.

## Acknowledgments

We gratefully thank Guiomar Wiesel for her technical assistance. We dedicate this work to Dr. Nilton Barreto dos Santos, a superb young scientist and a dear friend, victim of COVID-19.

**Supplementary information is available**

