## Supplemental Figures for "Genomic glucocorticoid receptor effects guide acute stress-induced delayed anxiety and basolateral amygdala spine plasticity in rats"

**This File includes:**

Figures S1 to S4

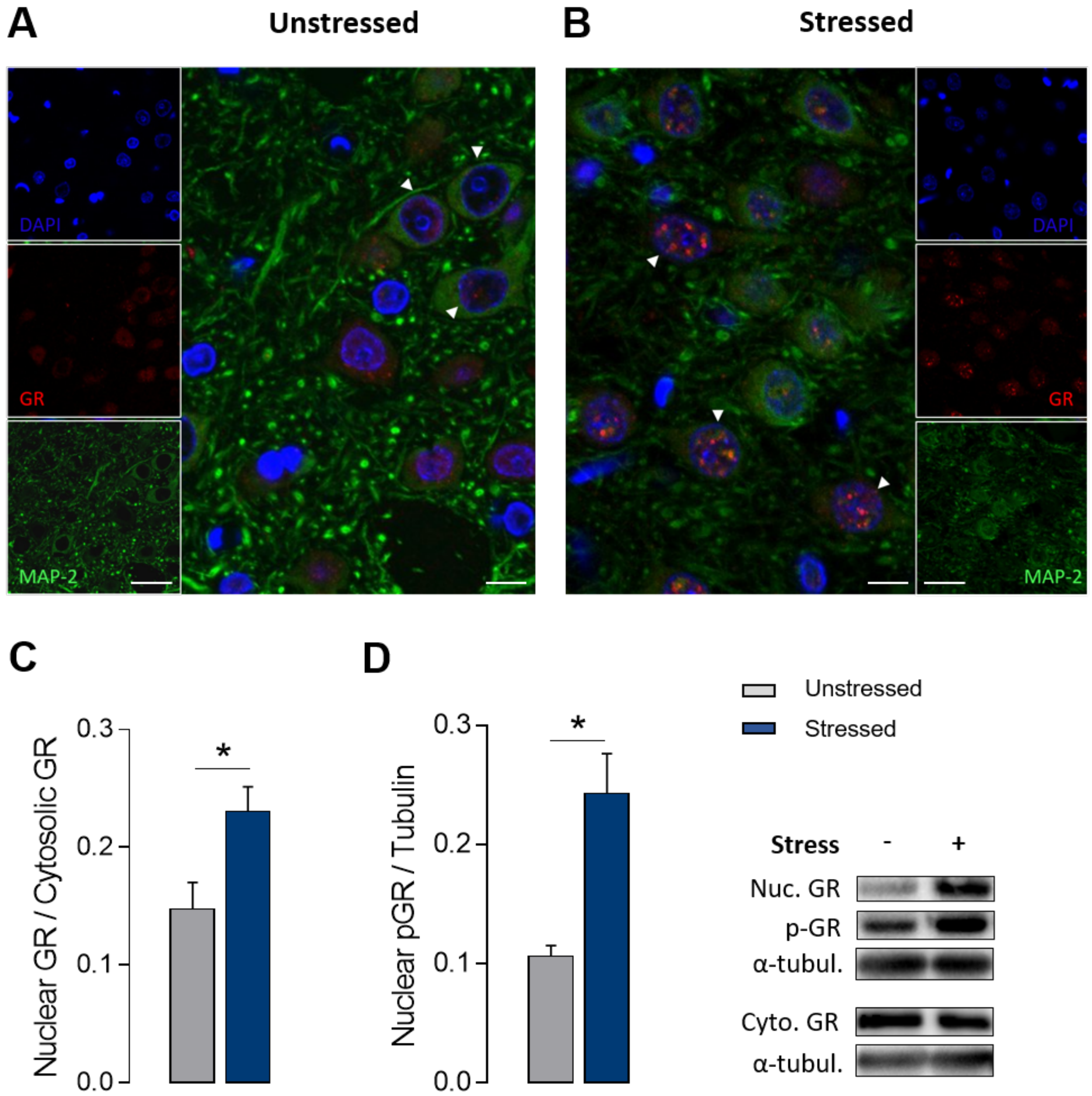


**Fig. S1.** Acute restraint stress effects on GR nuclear translocation and GRser211 phosphorylation in the BLA. (A and B) BLA immunofluorescence showing increased accumulation of GR (red) in the nucleus of neurons (green, MAP-2) from stressed animals (B) compared to non-stressed animals immediately after the cessation of stress (A). DAPI (blue) was used as a nuclear marker. Arrowheads indicate the nuclear region of neurons. (Scale bars on smaller and larger panels correspond to 25 µm and 8 µm, respectively). Images were acquired in a confocal microscope sectioned in the Z plane at 0.6 µm. (C and D) Western blot analysis showed an increase in GR nuclear translocation (C; n = 4-5 for each group, two-tailed student´s t-test: $P = 0.0346$), as well as in GRser211 phosphorylation (D; n = 3 for each group, two-tailed student´s t-test: $P = 0.0166$) in stressed animals compared to unstressed ones. (Bottom, right) Representative autoradiography of Western blot. Results are represented as mean ± SEM. Significance differences between groups are indicated as * $P < 0.05$.

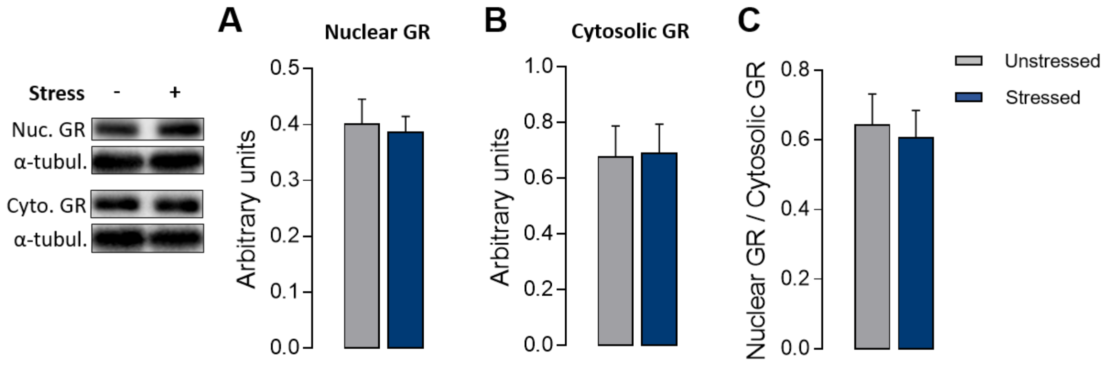


**Fig. S2.** GR expression in the BLA 10 days after the stress. There is no delayed effect of acute restraint stress on GR nuclear translocation in the BLA. (Left) Representative autoradiography of Western blot. (A–C) Western blot analysis showed no effect of stress on nuclear GR (A; n = 5 per group, two-tailed student´s t-test: P = 0.7837), cytosolic GR (B; n = 5 per group, two-tailed student´s t-test: P = 0.9408), or the nuclear-cytosolic ratio (C; n = 5 per group, two-tailed student´s t-test: P = 0.7571). Results are represented as mean ± SEM.

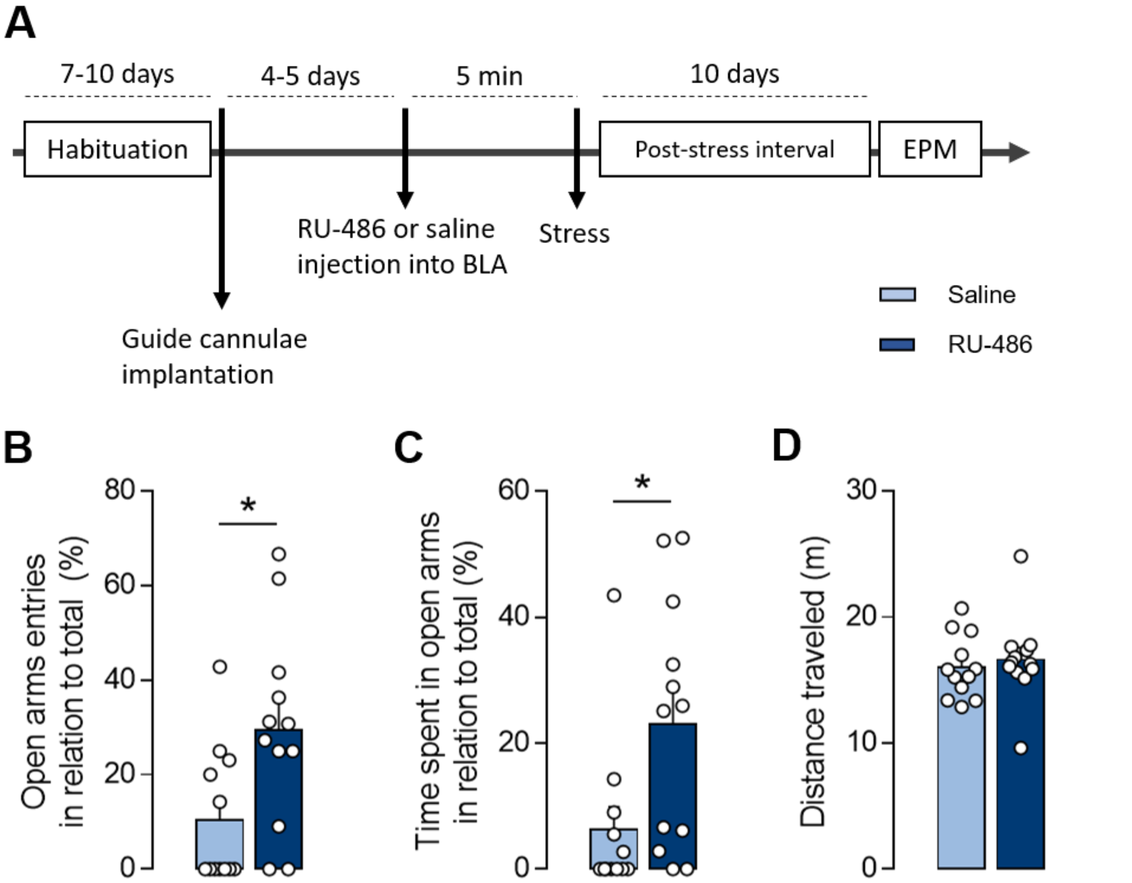


**Fig. S3.** Pre-stress intra-BLA GR antagonist RU-486 administration prevented the delayed emergence of stress-induced anxiety-like behavior. (A) Schematic representation of the experimental design showing the timeline of or RU-486 or saline injection into BLA, the stress submission, and the EPM behavioral test. (B–D) Pre-stress intra-BLA saline administered animals exhibited reduced open arms entries (B; n = 12 animal per group, two-tailed student´s t-test: P = 0.0161) and time spent (C; n = 12 animal per group, two-tailed student´s t-test: P = 0.0215) in the EPM test comparing to RU-486 administered animals. (D) There was no influence of intra-BLA RU-486 administration in the animal's locomotor activity in the EPM (n = 12 animal per group, two-tailed student´s t-test: P = 0.6168). Results are represented as mean ± SEM. Significance differences between groups are indicated as * P < 0.05.

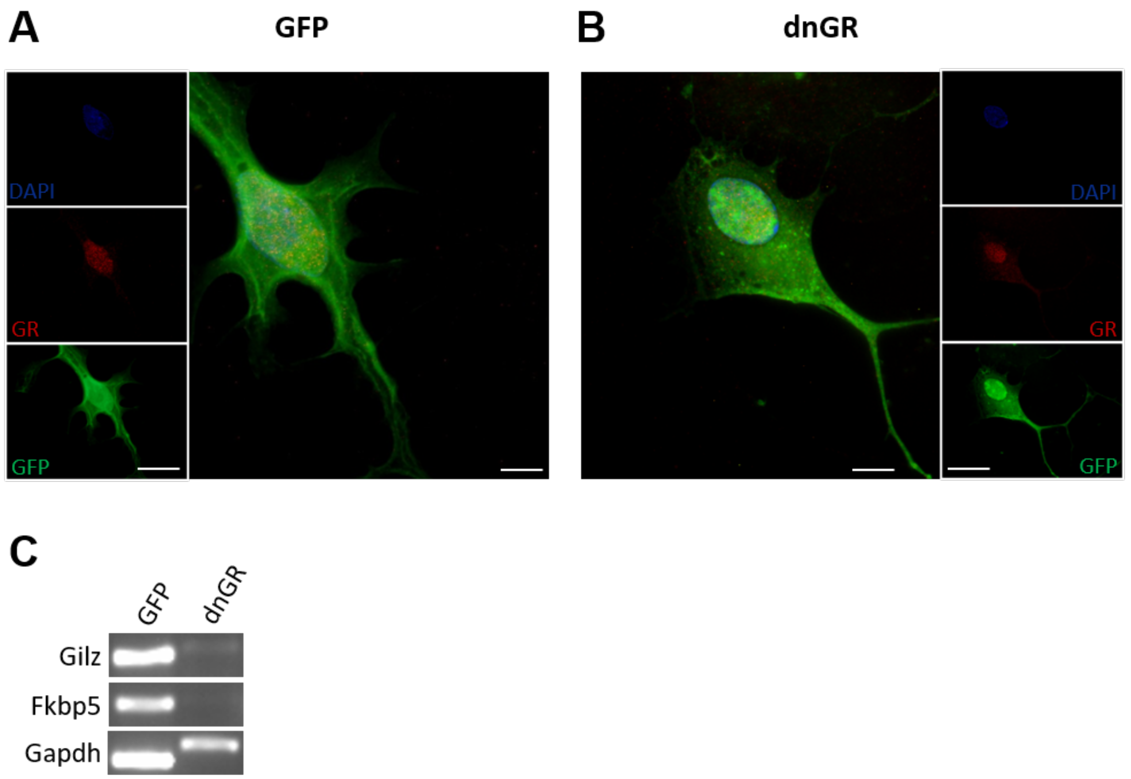


**Fig. S4.** dnGR- and GFP-infected cells response to the challenge with CORT. (A and B) Primary culture immunofluorescence of rat cortex showing GR (red) distribution in the cytoplasmic and nuclear compartments of neurons. Neurons were infected with GFP (A) or dnGR (B) and stimulated with CORT (10 μM, 1h). DAPI was used as a nuclear marker (Blue) and GFP is represented in green. (Scale bars on smaller and larger panels correspond to 30 μm and 10 μm, respectively). (C) RT-PCR cDNA amplicons for GILZ, FKBP5, and GAPDH in GFP- and dnGR-infected cells treated with CORT (10 μM, 1h).
